# Antioxidant role of the GABA shunt in regulating redox balance in blood progenitors during *Drosophila* hematopoiesis

**DOI:** 10.1101/2025.01.31.635852

**Authors:** Manisha Goyal, Sakshi Tiwari, Bruce Cooper, Ramaswamy Subramanian, Tina Mukherjee

## Abstract

Redox balance is crucial for normal development of stem and progenitor cells that reside in oxidative environments. This study explores the mechanisms of redox homeostasis in such niches and investigates myeloid-like blood progenitor cells that generate reactive oxygen species (ROS) and moderate it developmentally. Our findings reveal that during lymph gland development, as the blood-progenitor cells oxidize pyruvate via the TCA cycle leading to the generation of ROS, these cell also *de novo* synthesize GSH to counter excessive ROS and ensure redox balance. GABA metabolism, through GABA-shunt, restricts pyruvate dehydrogenase (PDH) activity and consequently TCA rate. This allows a metabolic route to sustain serine levels in these cells which is the rate limiting precursor controlling *de novo* GSH production. Disruption of GABA metabolism leads to a metabolic imbalance, characterized by excessive PDH activity, heightened TCA rate leading to impaired serine/GSH production and overall ROS dysregulation. Overall, the study presents a unique metabolic state whereby, in the blood-progenitor cells, by keeping PDH and TCA activity in check and promoting serine/GSH generation, GABA metabolism establishes a metabolic framework that optimises the use of ROS in blood-progenitors, while ensuring redox homeostasis.

## Introduction

Maintaining elevated reactive oxygen species (ROS) is a critical feature of many stem and progenitor cells (1,2). While the developmentally generated ROS is necessary for coordinating their normal homeostatic functions (3), the mechanisms by which cells thriving in oxidative environments prevent the accumulation of oxidative stress remains an active area of research.

Specifically, blood stem and progenitor cells of common myeloid origin rely on mitochondrially generated ROS for their development (4). However, any processes that excessively increases mitochondrial oxidation or disrupts mitochondrial function leading to aberrant ROS generation can impair progenitor maintenance and result in differentiation defects (5,6). Therefore, it is crucial to maintain ROS levels within a specific threshold that allows ROS to function as a signaling molecule rather than a stress component. In this regard, the role of ROS scavenging and antioxidant mechanisms, which restore balance when ROS levels are excessively high, becomes increasingly relevant (7–9). These mechanisms include both enzymatic and non-enzymatic processes, with antioxidant enzymes such as peroxidase, catalase, and superoxide dismutase (SOD), along with non-enzymatic metabolites like ascorbate, glutathione (GSH) and vitamin A, and form an integral part of the ROS scavenging system (10). The generation and scavenging of ROS is shaped by various cellular changes (11). In this study, we examine how hematopoietic progenitor cells, which develop in a dynamic niche, and are influenced by a range of environmental influences (12–16), coordinate ROS production and scavenging processes, so as to maintain redox balance.

Lymph gland, the primary definitive hematopoietic organ of *Drosophila*, present during larval stages of development, is a multilobed structure that houses multipotent stem-like blood progenitor cells, which resemble vertebrate myeloid cells (reviewed in (17). These progenitors reside in the inner core region of the organ termed Medullary Zone (MZ) and differentiate into all mature blood cell types, which are then localized in the outer zone of the lymph gland termed Cortical Zone (CZ) (17,18). Similar to common myeloid progenitor cells (19), lymph gland progenitors have been shown to maintain higher ROS compared to their surrounding differentiated cells. In these progenitor cells, the developmentally generated ROS is essential for priming the cells toward differentiation cues (5). Various signaling cues including NF-κB (20,21), JNK (22), Notch (23) and cues of nutritional/metabolic in origin like fatty acids (24) have been shown to regulate blood progenitor development by modulating ROS levels in the lymph gland (2). However, any aberrant ROS levels impairs progenitor development and homeostasis, thus highlighting the importance of ROS regulation in progenitor development (5). ROS generation in progenitor cells is linked to mitochondrial/TCA activity (5,25), and antioxidants such as catalase, SOD, and glutathione peroxidase (Gtpx) have been shown to play essential roles in maintaining ROS homeostasis (5). These antioxidants help neutralize excess ROS, protecting progenitor cells from oxidative damage and ensuring proper development and function. Therefore, the balance between ROS production and antioxidant activity is essential for controlled differentiation and survival of these myeloid-like blood progenitor cells. The understanding of regulatory mechanisms that coordinate ROS generation with scavenging to maintain a delicate ROS balance in the lymph gland blood system forms the central question of the current study.

Our previous research identified the tricarboxylic acid (TCA) cycle as a source of ROS in the progenitor cells (25). Moreover, we found that neuronally derived GABA (γ-aminobutyric acid, (13), its uptake and metabolism by the progenitor cells (26) control ROS production through limiting TCA cycle activity (25). The study demonstrated that GABA taken up by progenitor cells via the GABA transporter (Gat) and metabolized through the GABA-shunt pathway, produces succinate, which regulates pyruvate dehydrogenase (PDH) activity and controls the rate of the TCA cycle (Fig. 1A). Loss of GABA function led to increased TCA activity and elevated ROS levels in the progenitor cells and highlighted a key redox modulatory role for GABA in the progenitor cells.

**Figure 1.**
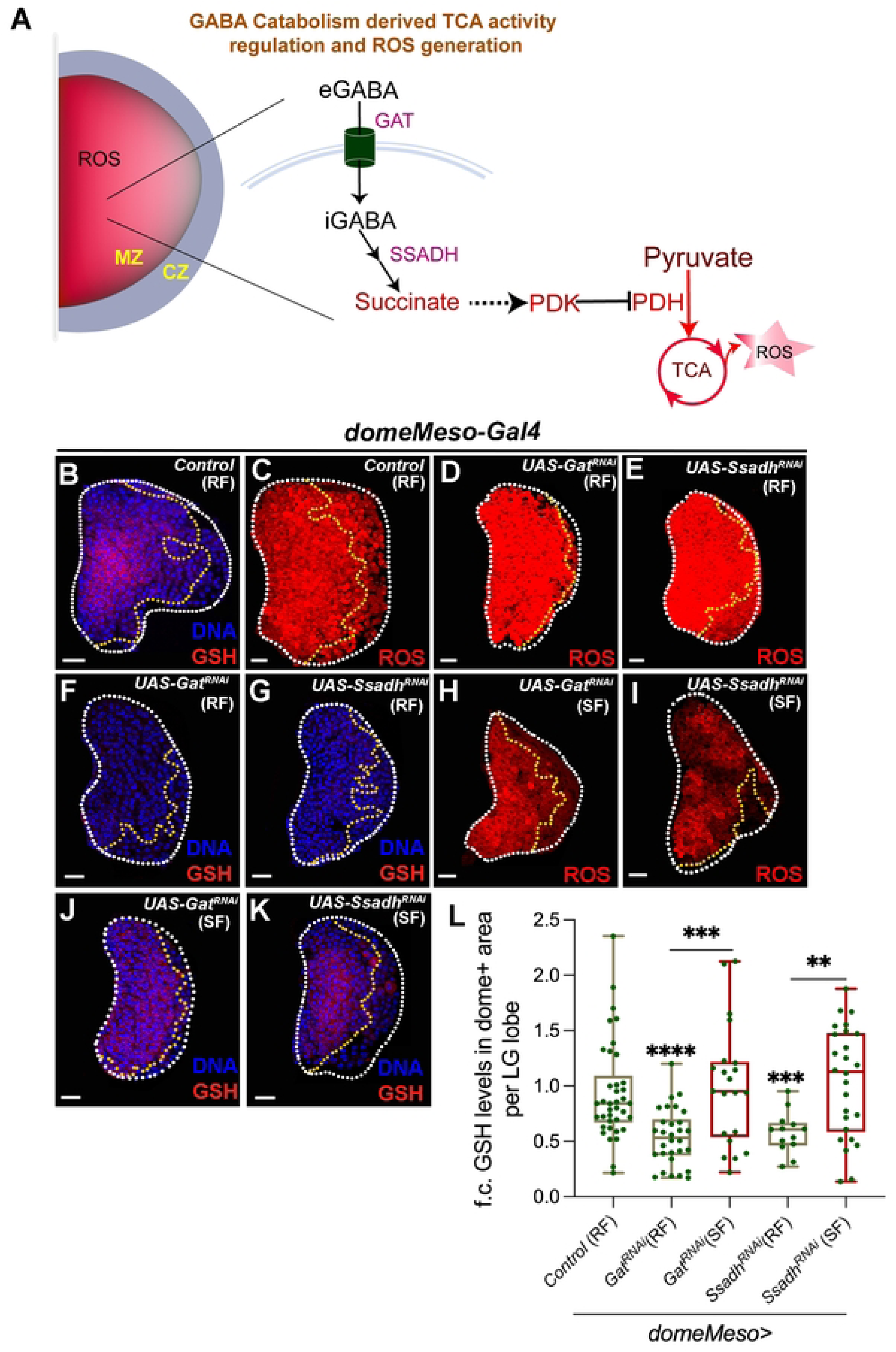
GABA catabolic pathway in *Drosophila* blood-progenitor cells control their GSH levels. RF is regular food; SF is succinate food. Data is presented as median plots (**p<0.01; ***p<0.001 and ****p<0.0001). Mann-Whitney test is applied for **L**. In **L**, ‘n’ is total number of lymph gland lobes analysed and in the graph each lobe is represented by a green dot. Scale bar: 20µm. DNA is stained with DAPI in blue, comparisons for significance are done with their respective control and also with respective genetic conditions which are indicated by horizontal lines drawn above the boxplots. Red bars represent rescue combinations. White border demarcates the lymph gland lobe and yellow border marks the dome positive area towards the left side and the respective lymph gland images with dome^+^ (green) is shown in **Fig. S1**. MZ-medullary zone (marked in green), CZ-cortical zone (marked in blue), ROS-reactive oxygen species, GABA-γ-aminobutyric acid, eGABA-extracellular GABA, iGABA-intracellular GABA, SSADH-succinic semialdehyde dehydrogenase, PDK-pyruvate dehydrogenase kinase, PDH-pyruvate dehydrogenase, TCA-tricarboxylic acid. **(A)** Schematic representation of GABA catabolism and its regulation of TCA activity. In blood-progenitor cells (MZ), which maintain elevated levels of ROS, extracellular GABA (eGABA) is internalized by GABA transporter (GAT), that is further catabolized into succinate by a two-step process including second, rate-limiting step catalysed by SSADH. GABA catabolism derived succinate regulates PDK activity (pPDK) to control PDH phosphorylation, which restricts TCA cycle activity and generation of ROS. **(B-G)** Representative images showing ROS (red) and GSH (red) levels in the lymph gland progenitor cells (area marked within the yellow dotted line) from different genetic backgrounds. Control (*domeMeso-Gal4,UAS-GFP/+*) showing **(B)** GSH levels, which is elevated in the progenitor cells and **(C)** elevated ROS (stained with DHE), in them. Progenitor specific expression of **(D, F)** *Gat^RNAi^* (*domeMeso-Gal4,UAS-GFP;UAS-Gat^RNAi^*) and **(E, G)** *Ssadh^RNAi^* (*domeMeso-Gal4,UAS-GFP;UAS-Ssadh^RNAi^*) leads to an increase in their progenitor (**D, E**) ROS levels and reduction in **(F, G)** GSH. For comparison, see control ROS in **(C)** and control GSH (B). For quantifications, refer to **L**. (**H-K)** Succinate supplementation of **(H, J)** *Gat^RNAi^* (SF, *domeMeso-Gal4,UAS-GFP;UAS-Gat^RNAi^*) and **(I, K)** *Ssadh^RNAi^* (SF, *domeMeso-Gal4,UAS-GFP;UAS-Ssadh^RNAi^*) larvae, rescues blood-progenitor **(H, I)** ROS and **(J, K)** GSH levels respectively. For comparison, see larvae raised on regular food of *Gat^RNAi^* **(D, F)** and *Ssadh^RNAi^* **(E, G)** and control in **B**. For quantifications, refer to **L**. **(L)** Quantification of blood-progenitor GSH levels (fold change, f.c.) in *domeMeso>GFP/+* (control, n=38)*, domeMeso>GFP/Gat^RNAi^* (RF, n=30, p<0.0001), *domeMeso>GFP/Gat^RNAi^* (SF, n=21, p=0.0009, in comparison to *Gat^RNAi^*), *domeMeso>GFP/Ssadh^RNAi^* (RF, n=13, p=0.0007) and *domeMeso>GFP/ Ssadh^RNAi^* (SF, n=26, p=0.0053, in comparison to *Ssadh^RNAi^*).

The findings from our previous work, led us to further investigate the redox modulatory effects of the GABA-shunt pathway. In this study, we describe its role in the generation of a key antioxidant, glutathione (GSH), by the blood-progenitor cells and establishment of their anti-oxidant potential. Glutathione serves as a substrate for enzymes like glutathione S-transferase, and Gtpx, which are essential for mitigating redox stress and preventing oxidative damage (27,28). These enzymatic ROS scavengers have been shown to play a vital role in the maintenance of the lymph gland progenitor cells in their undifferentiated state (5). Thus the heightened reliance of progenitor cells on these enzymatic scavengers led us to explore, glutathione regulation and we observed that progenitor cells maintained elevated levels of GSH in them. The accumulation of GSH in progenitor cells in addition to elevated ROS also detected in them, suggested specialized developmental program that operated towards maintaining effective ROS scavenging mechanisms in the myeloid-like blood progenitor cells.

Glutathione is a tripeptide comprised of three amino acids (cysteine, glutamate, and glycine), and cycles between oxidized and reduced states (29). The presence of total GSH in the progenitor cells alluded to its *de novo* generation. By employing a combination of immunohistochemical and mass spectrometry-based approaches, we investigated the underlying modality leading to GSH accumulation in progenitor cells. The investigations revealed an unexpected control of pyruvate metabolism in establishing GSH levels in them. Active pyruvate oxidation and TCA cycling generates ROS in progenitor cells, but the extent of pyruvate utilization and TCA activity defines the cellular potential to generate serine. Serine is a key biosynthetic intermediate to generate GSH, and its levels become rate-limiting in defining progenitor GSH levels. Thus, to achieve this balance, GABA metabolism coordinates pyruvate utilization in the progenitor cells whereby it allows controlled pyruvate entry into the TCA and simultaneously enables a state that sustains serine synthesis. We speculate that GABA metabolism diverts pyruvate to generate serine via gluconeogenesis, and together this ascertains a metabolic program that allows progenitor cells to develop homeostatically, producing ROS while simultaneously scavenging it. The use of GABA metabolism by the blood progenitor cells to counter redox stress at the nexus of regulating pyruvate metabolism is a central finding from the study. The reliance of vertebrate hematopoietic stem and progenitor cells on TCA/mitochondria to generate ROS (30–33) and their dependency on GABA is much recently shown (34). Therefore, we speculate that the metabolic program identified in this study may well be conserved in the mammalian hematopoietic niche and also across other stem and progenitor cells residing in oxidative niches (19,35–37).

## Results

### Control of blood progenitor glutathione levels by the GABA-shunt pathway

The non-enzymatic antioxidant glutathione (GSH) was analyzed in *Drosophila* third instar larval lymph glands using a specific antibody generated against it (abcam 9443, (38). The results revealed that GSH levels were significantly higher in progenitor cells in comparison to the differentiated cells of the cortical zone (CZ) (Fig. 1B and Fig. S1D, D’). Notably, the antibody detects total glutathione and does not distinguish between its oxidized and reduced forms (38). As a proxy for the specificity of the antibody against total GSH, we reared control larvae from early first instar developmental stage on food supplemented with 0.1% GSH and analyzed these lymph glands for their GSH levels. Wandering 3^rd^ instar larvae obtained from this condition showed increased progenitor GSH levels (Fig. S1A-C), and implied the antibody detected total GSH. The elevated GSH detected in progenitor cells was also consistent with enzymatic antioxidant activities linked to glutathione reported in literature. Specifically, glutathione S-transferase activity, observed with *gstD-GFP* reporter, shows heightened expression in the MZ progenitor cells (20) and secondly, glutathione peroxidase (Gtpx), whose function is necessary for progenitor homeostasis (5). Together with these previously reported findings, the elevated GSH data corroborated with its requirement in the progenitor cells and implicated regulatory mechanisms governing GSH homeostasis in them, which we set out to investigate.

We have previously demonstrated that GABA metabolism is crucial for moderating progenitor TCA activity and plays an important role in regulating ROS generation. Furthermore, ROS moderation through GABA catabolism via controlling TCA activity is central to regulate homeostatic lymph gland growth (25). We found that down-regulating components of the GABA catabolic pathway, not only affected progenitor ROS levels (Fig. 1C-E and Fig. S1E-G) but also led to reduction in their GSH levels (Fig. 1F, G, L and Fig. S1H-I’). Specifically, blocking progenitor GABA uptake, by knocking down the transporter, *Gat* (Fig. 1F and Fig. S1H, H’) or downregulating the rate-limiting GABA-shunt catabolic enzyme, *Ssadh* (Fig. 1G and Fig. S1I, I’), using the progenitor-specific driver *domeMeso-Gal4,* resulted in a significant reduction in their GSH levels. Succinate, which is the end-product of GABA breakdown, supplementing it in *Gat* or *Ssadh* RNA*i* expressing conditions, that restores progenitor ROS levels (Fig. 1H, I and Fig. S1J, K), also caused a significant recovery of progenitor GSH levels (Fig. 1J, K, L and Fig. S1L-M’). These data suggested an unexpected role for GABA shunt pathway in progenitor GSH homeostasis alongside its known role in moderating ROS generation that we have previously reported. The mechanism by which GABA influenced GSH levels, was therefore explored.

### Regulation of glutathione synthesis by GABA catabolic pathway

The antibody used to detect GSH does not distinguish between its oxidized and reduced forms; therefore, the measured GSH levels in progenitor cells represented its total concentration (38). The observed reduction in GSH levels in *Gat* and *Ssadh* mutants, followed by recovery upon succinate supplementation (Fig. 1), led us to hypothesize a defect in GSH synthesis, as opposed to any changes in its oxidized or reduced state. This prompted us to explore the mechanisms regulating GSH synthesis and the involvement of GABA metabolism in this process.

Glutathione (GSH) is a non-protein thiol metabolite synthesized from the amino acid cysteine, glutamate, and glycine (Fig. 2A). The rate-limiting step in GSH synthesis involves the combination of cysteine and glutamate to form γ-glutamyl cysteine. In the final step, glycine is added, resulting in the formation of GSH (Fig. 2A) (29). To determine whether the reduction observed in GABA catabolic mutants was due to changes in the intermediates required for GSH synthesis, we analyzed progenitor levels of these intermediates using specific antibodies, where available and also via mass spectrometry (see Methods).

**Figure 2.**
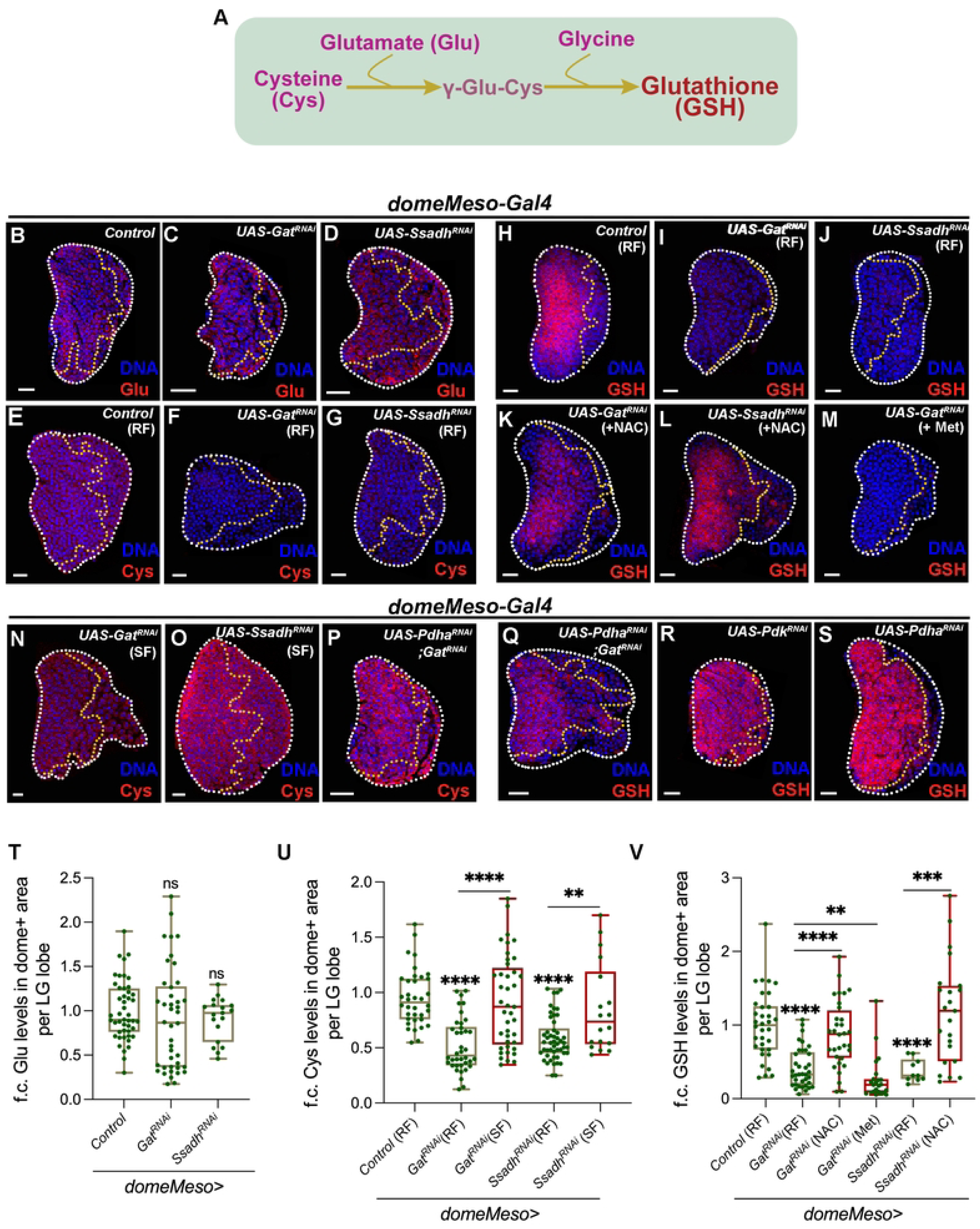
Regulation of glutathione synthesis by GABA catabolic pathway. RF is regular food, SF is succinate food, NAC is N-acetyl cysteine and Met is methionine supplemented food. Data is presented as median plots (**p<0.01; ***p<0.001; ****p<0.0001 and ns is non-significant). Mann-Whitney test is applied for **T**-**V**. In **T-V**, ‘n’ is total number of lymph gland lobes analysed and in the graph each lobe is represented by a green dot. Scale bar: 20µm. DNA is stained with DAPI in blue, comparisons for significance are done with their respective control and also with respective genetic conditions which are indicated by horizontal lines drawn above the boxplots. Red bars represent rescue combinations. White border demarcates the lymph gland lobe and yellow border marks the dome positive area towards the left side and the respective lymph gland images with dome^+^ (green) is shown in **Fig. S2**. **(A)** Schematic representation of glutathione (GSH) synthesis pathway. Cysteine (Cys) and glutamate (Glu) conjugate to form γ-glutamyl-cysteine, and further addition of glycine leads to GSH formation. **(B-D)** Representative images showing glutamate (Glu, red) levels in lymph gland progenitor cells across different genetic backgrounds. In comparison to **(B)** control (*domeMeso-Gal4,UAS-GFP/+*) lymph gland showing uniform glutamate levels across all cells of the tissue, including the progenitor-cells (area within the yellow dotted line). Expressing **(C)** *Gat^RNAi^* (*domeMeso-Gal4,UAS-GFP;UAS-Gat^RNAi^*) and **(D)** *Ssadh^RNAi^* (*domeMeso-Gal4,UAS-GFP;UAS-Ssadh^RNAi^*) in the progenitor cells does not affect their glutamate levels. For quantifications, refer to **T**. **(E-G)** Representative images showing cysteine (Cys, red) expression in lymph gland progenitor cells (area marked within the yellow dotted line) from different genetic backgrounds. **(E)** Control (RF, *domeMeso-Gal4,UAS-GFP/+*) lymph gland showing relatively uniform cysteine levels in all cells of the lymph gland including progenitor-cells (area demarcated within the yellow border). Expressing **(F)** *Gat^RNAi^* (RF, *domeMeso-Gal4,UAS-GFP;UAS-Gat^RNAi^*) and **(G)** *Ssadh^RNAi^* (RF, *domeMeso-Gal4,UAS-GFP;UAS-Ssadh^RNAi^*) in the progenitor cells leads to reduction in cysteine levels as compared to control **(E)**. For quantifications, refer to **U**. **(H-M)** Representative images showing GSH (red) expression in lymph gland progenitor cells (area marked within the yellow dotted line) from different genetic backgrounds. **(H)** Control (RF, *domeMeso-Gal4,UAS-GFP/+*) lymph gland showing GSH levels. While expressing **(I)** *Gat^RNAi^* (RF, *domeMeso-Gal4,UAS-GFP;UAS-Gat^RNAi^*) and **(J)** *Ssadh^RNAi^* (RF, *domeMeso-Gal4,UAS-GFP;UAS-Ssadh^RNAi^*) in the progenitor cells leads to reduction in GSH levels, supplementing these genetic conditions with **(K, L)** N-acetyl cysteine (NAC), restores GSH levels in both **(K)** *Gat^RNAi^* (NAC, *domeMeso-Gal4,UAS-GFP;UAS-Gat^RNAi^*) and **(L)** *Ssadh^RNAi^* (NAC, *domeMeso-Gal4,UAS-GFP;UAS-Ssadh^RNAi^*) background to levels almost comparable to control shown in **H**. **(M)** Methionine supplementation to *Gat^RNAi^* (Met, *domeMeso-Gal4,UAS-GFP;UAS-Gat^RNAi^*) does not recover GSH levels. For comparison also refer to *Gat^RNAi^* **(I)** and *Ssadh^RNAi^* **(J)** raised on regular food (RF). For quantifications, refer to **V**. **(N-O)** Succinate supplementation to **(N)** *Gat^RNAi^* (SF, *domeMeso-Gal4,UAS-GFP;UAS-Gat^RNAi^*) and **(O)** *Ssadh^RNAi^* (SF, *domeMeso-Gal4,UAS-GFP;UAS-Ssadh^RNAi^*) restores blood-progenitor cysteine levels. For comparison see lymph gland lobes from **(F)** *Gat^RNAi^* and **(G)** *Ssadh^RNAi^* animals raised on regular food (RF) and control in **E**. For quantifications, refer to **U**. **(P-Q)** Progenitor specific expression of *Pdha^RNAi^* in *Gat^RNAi^* animals (*domeMeso-Gal4,UAS-GFP; UAS-Pdha^RNAi^;UAS-Gat^RNAi^*) leads to a recovery of blood-progenitor **(P)** cysteine and **(Q)** GSH levels. Compare with **(F)** Cys and **(I)** GSH in *Gat^RNAi^ (domeMeso-Gal4,UAS-GFP;UAS-Gat^RNAi^*). For quantifications, refer to **Fig. S2V** and **Fig. S2W**. **(R, S)** Expressing **(R)** *Pdk^RNAi^* (*domeMeso-Gal4,UAS-GFP;UAS-Pdk^RNAi^*) or **(S)** *Pdha^RNAi^* (*domeMeso-Gal4,UAS-GFP;UAS-Pdha^RNAi^*), specifically in the progenitor cells, does not reveal any dramatic change in GSH levels. Compare to control **(H).** For quantifications, refer to **Fig. S2W**. **(T)** Quantification of blood-progenitor glutamate (Glu) levels (fold change, f.c.) in *domeMeso>GFP/+* (control, n=47)*, domeMeso>GFP/Gat^RNAi^* (n=42, p=0.1570) and *domeMeso>GFP/Ssadh^RNAi^* (n=19, p=0.5180). **(U)** Quantification of blood-progenitor cysteine (Cys) levels (fold change, f.c.) in *domeMeso>GFP/+* (control, RF, n=36)*, domeMeso>GFP/Gat^RNAi^* (RF, n=38, p<0.0001), *domeMeso>GFP/Gat^RNAi^* (SF, n=41, p<0.0001 in comparison to *Gat^RNAi^*, RF), *domeMeso>GFP/Ssadh^RNAi^* (RF, n=50, p<0.0001), and *domeMeso>GFP/Ssadh^RNAi^* (SF, n=18, p=0.0075 in comparison to *Ssadh^RNAi^* on RF). **(V)** Quantification of blood-progenitor GSH levels (fold change, f.c.) in *domeMeso>GFP/+* (control, RF, n=34)*, domeMeso>GFP/Gat^RNAi^* (RF, n=37, p<0.0001), *domeMeso>GFP/Gat^RNAi^* (NAC, n=32, p<0.0001 in comparison to *Gat^RNAi^*, RF), *domeMeso>GFP/Gat^RNAi^* (Methionine, n=23, p=0.0045 in comparison to *Gat^RNAi^*, RF), *domeMeso>GFP/Ssadh^RNAi^* (RF, n=10, p<0.0001), and *domeMeso>GFP/Ssadh^RNAi^* (NAC, n=23, p=0.0006 in comparison to *Ssadh^RNAi^*, RF).

We found that glutamate levels upon disruption of the *GABA transporter (Gat)* or *Ssadh* in blood progenitor cells revealed no significant change both immunohistochemically with an anti-glutamate antibody (Fig. 2B-D, T and Fig. S2A-C) and mass-spectrometrically (Fig. S2T). However, the levels of cysteine when assessed using a cysteine-specific antibody, showed a reduction in its levels (Fig. 2E-G, U). In control lymph glands, cysteine is detected uniformly across progenitor and differentiating cells (Fig. 2E and Fig. S2D), however with progenitor-specific loss of *Gat* or *Ssadh*, it led to a significant reduction in overall cysteine levels (Fig. 2E-G, U and Fig. S2D-F). Cysteine levels were also analyzed mass-spectrometrically, but we failed to detect any signal and speculate this to be due to limited sample size and cysteine’s inherent instability due to its thiol group (39). Glycine was assessed using mass-spectrometry and using this approach we observed no detectable changes for glycine levels in *Gat* and *Ssadh* loss of function condition (Fig. S2U). These data revealed that GABA catabolism most likely by controlling progenitor cysteine levels controlled their GSH production. The reduction in GSH in these catabolic mutants could therefore arise as a consequence of reduced cysteine levels.

Therefore, to address if the lower amounts of cysteine led to reduction in GSH synthesis, we supplemented *domeMeso>Gat^RNAi^* and *domeMeso>Ssadh^RNAi^* conditions with N-acetylcysteine (NAC), an analog for cysteine. This supplementation restored progenitor GSH levels in *Gat^RNAi^* and *Ssadh^RNAi^* to nearly comparable levels seen in control lymph glands (Fig. 2H-L, V and Fig. S2G-K). Importantly, cysteine is a sulfur amino acid and can be derived from methionine through its cycling and the transsulfuration pathway (40). Therefore, we supplemented *Gat^RNAi^* mutant larvae with methionine, however, its supplementation failed to show any restoration of GSH levels (Fig. 2M, V and Fig. S2L). This data indicated that NAC mediated recovery of GSH was specific to restoration of cysteine and not a non-specific rescue by any other sulfur amino acid. Furthermore, the data also suggested that the cysteine synthesis regulation by GABA catabolism was independent of its regulation of methionine. Therefore, we conducted the next set of analyses to investigate how GABA catabolism supported cysteine synthesis in progenitor cells.

We know that downstream of GABA catabolism, the succinate derived from its breakdown restored GSH levels in the mutant backgrounds (Fig. 1L). If succinate supplementation in *domeMeso>Gat^RNAi^* and *domeMeso>Ssadh^RNAi^* larvae effectively restored cysteine levels was therefore asked. Interestingly, we observed that succinate restoration could re-establish cysteine in the lymph glands (Fig. 2N, O, U and Fig. S2M, N). These results suggested that GABA catabolism derived succinate controls progenitor cysteine levels.

### Pyruvate metabolism and its control on progenitor GSH generation

Our previous work has shown that, GABA-shunt derived succinate moderates PDH activity to control the entry of pyruvate into the TCA cycle, resulting in restricted TCA rate and consequently ROS generation (25). If elevated PDH activity in loss of GABA metabolism contributed to any lowering of GSH synthesis was therefore explored. For this, we down-regulated *Pdha* in *Gat^RNAi^* condition and assessed for cysteine and GSH levels. We found that this genetic condition restored lymph cysteine levels, almost comparable to that seen in control lymph glands (Fig. 2P and Fig. S2O, V) and very importantly progenitor GSH levels were restored as well (Fig. 2Q and Fig. S2P, W). The data implicated elevated PDH activity in absence of GABA metabolism as the underlying cause for dampened blood progenitor GSH production as well. The regulation of pyruvate oxidation by PDH has emerged as a necessary step not only for maintaining TCA rate but also for supporting the antioxidant potential i.e. by driving cysteine and consequently GSH synthesis in the progenitor cells.

Therefore, to address the influence of pyruvate oxidation in progenitor GSH levels and if this was purely maintained as a consequence of PDH activity, we increased PDH activity independent of altering GABA catabolism in the progenitor cells. We undertook this by downregulating *Pdk*, the kinase that phosphorylates PDH and inactivates it (41,42). *Pdk* downregulation in progenitor cells did not affect their GSH levels (Fig. 2R and Fig. S2Q, W). The data was intriguing and showed that heightened pyruvate oxidation via increasing PDH activity was insufficient to impair GSH generation, even though we knew this genetic condition could increase progenitor ROS levels (25). Contrary to PDH activation, we also dampened it and blocked pyruvate oxidation by down-regulating *Pdha* in blood-progenitors. This condition as well did not reveal any major difference in progenitor GSH or cysteine levels (Fig. 2S and Fig. S2R, S-S’, V, W). Thus, the data suggested that moderating PDH function alone did not influence GSH synthesis and that GABA catabolism influenced additional aspects of pyruvate metabolism to control Cys/GSH synthesis in progenitor cells. The down-regulation of *Pdha* in *Gat^RNAi^* most likely restored pyruvate levels in these cells which consequently enabled their cysteine and GSH synthesis. In this regard, pyruvate and its availability emerged as a key cellular metabolite in governing progenitor Cys and GSH levels.

Pyruvate is an essential cellular metabolite and plays not just an important role in driving the TCA (43) but also a variety of other intracellular processes, like glycolysis, gluconeogenesis and fatty acid synthesis (44). Pyruvate’s utilization as a gluconeogenic intermediate is key for driving generation of metabolites, including amino acids and nucleotides (45,46). Specific to blood progenitor cells, apart from its oxidation to drive progenitor ROS generation, our previous studies have highlighted its importance in promoting a glycolytic state in progenitor cells, where the stabilization of Hif-1α (Sima) by the GABA shunt pathway reinforces their glycolytic potential (26). Thus, controlling pyruvate metabolism in blood progenitor may be central to facilitate its diverse metabolic outcomes. In this regard, the importance of GABA-shunt and its regulatory role on influencing pyruvate metabolism to systematically enable cysteine and GSH synthesis was proposed.

### GABA catabolism controls progenitor cysteine synthesis via regulating de novo serine synthesis

This initiates an investigation into the mechanisms that lead to cysteine generation in blood-progenitor cells and any underlying connection to pyruvate metabolism.

Cysteine is a non-essential amino acid and is synthesized in cells via the conserved trans-sulfuration pathway (TSP) (Fig. 3A). Methionine and serine, the precursor amino acids in TSP, combines through a series of steps to gives rise to cysteine (40). While methionine supplementation data shown above (Fig. 2M) ruled out any contribution of it in GSH synthesis, any limitation in serine levels culminating in reduced cysteine generation was investigated. Serine is a component of TSP and when analysed for total serine in *Gat^RNAi^* conditions, we observed a significant decrease in total lymph gland serine levels (Fig. 3B). If the reduction in serine contributed towards cysteine and GSH generation in *Gat^RNAi^* condition was next assessed. We undertook supplementation of serine in *Gat^RNAi^* mutants and observed a significant rescue in both cysteine (Fig. 3C-E, L and Fig. S3A-C) and GSH (Fig. 3F-H, M and Fig. S3D-F) levels. Moreover, supplementing serine also restored the increased ROS phenotype detected in *Gat^RNAi^* (Fig. 3I-K’, N and Fig. S3G-I) condition to moderate levels. Consistent with lymph gland growth inhibitory effect of heightened ROS production (25), the small sized lymph gland phenotype in *Gat^RNAi^* condition was also restored by serine supplementation to sizes almost comparable to control lymph glands (Fig. 3K, K’ and O). These data implied that GABA catabolism in lymph gland blood progenitor cells controlled their serine levels, whose availability, through GSH generation controlled progenitor ROS scavenging. The dual control exerted by GABA catabolism on ROS production and progenitor ROS scavenging overall controlled homeostatic lymph gland growth and development.

**Figure 3.**
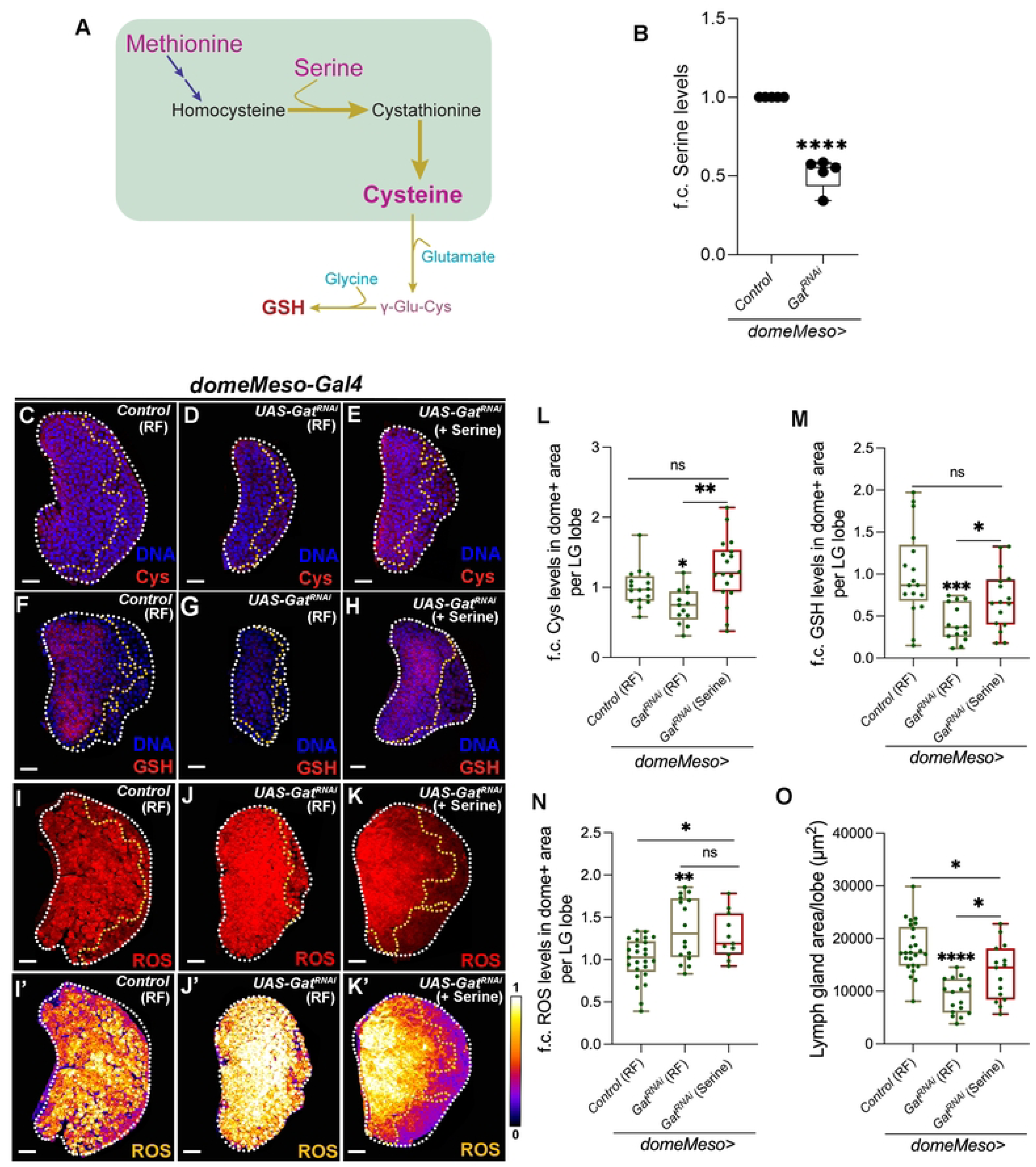
GABA catabolism controls progenitor cysteine synthesis via regulating *de novo* serine synthesis and maintain ROS homeostasis in the progenitor cells. RF is regular food, and serine is serine supplemented food. Data is presented as median plots (*p<0.05; **p<0.01; ***p<0.001; ****p<0.0001 and ns is non-significant). Two-way ANOVA is applied for **B** and Mann-Whitney test is applied for **L-O**. In **B** ‘N’ is number of experimental repeats and ‘n’ is sample size and shown by black dot (N). In **L-O**, ‘n’ is total number of lymph gland lobes analysed and in the graph each lobe is represented by a green dot. Scale bar: 20µm. DNA is stained with DAPI in blue, comparisons for significance are done with their respective control and also with respective genetic conditions which are indicated by horizontal lines drawn above the boxplots. Red bars represent rescue combinations. White border demarcates the lymph gland lobe and yellow border marks the dome positive area towards the left side and the respective lymph gland images with dome^+^ (green) is shown in **Fig. S3**. **(A)** Schematic representation of transsulfuration pathway (shown in the green box) and GSH synthesis. Methionine and serine combine to form cysteine via the transsulfuration pathway (TSP), which further combines with glutamate and glycine to form GSH. **(B)** Relative steady state levels (fold change, f.c.) of serine in *domeMeso>GFP/+* (control, N=5, n=15) *and domeMeso>GFP/Gat^RNAi^* (N=5, n=18, p<0.0001), where *Gat^RNAi^* in the progenitor cells leads to reduction in lymph gland serine levels as compared to control. **(C-E)** Representative images showing cysteine (Cys, red) expression in lymph gland progenitor cells (area marked within the yellow dotted line) from different genetic backgrounds. **(C)** Control (RF, *domeMeso-Gal4,UAS-GFP/+*) lymph gland showing relatively uniform cysteine levels in all cells of the lymph gland including progenitor-cells (area demarcated within the yellow border). While, expressing **(D)** *Gat^RNAi^* (RF, *domeMeso-Gal4,UAS-GFP;UAS-Gat^RNAi^*) in the progenitor cells leads to reduction in cysteine levels as compared to control **(C)**, supplementing this genetic condition with **(E)** serine (Serine, *domeMeso-Gal4,UAS-GFP;UAS-Gat^RNAi^*) recovers cysteine levels almost comparable to control **(C)**. For comparison, also refer to **(D)** *Gat^RNAi^* raised on regular food (RF). For quantifications, refer to **L**. **(F-H)** Representative images showing GSH (red) expression in lymph gland progenitor cells (area marked within the yellow dotted line) from different genetic backgrounds. **(F)** Control (RF, *domeMeso-Gal4,UAS-GFP/+*) lymph gland showing GSH levels in lymph gland progenitor-cells (area demarcated within the yellow border). While, expressing **(G)** *Gat^RNAi^* (RF, *domeMeso-Gal4,UAS-GFP;UAS-Gat^RNAi^*) in the progenitor cells leads to reduction in GSH levels as compared to control **(F)**, supplementing this genetic condition with **(H)** serine (Serine, *domeMeso-Gal4,UAS-GFP;UAS-Gat^RNAi^*) recovers GSH levels almost comparable to control **(F)**. For comparison, also refer to **(G)** *Gat^RNAi^* raised on regular food (RF). For quantifications, refer to **M**. **(I-K’)** Representative images showing ROS levels (area marked within the yellow dotted line) from different genetic backgrounds, shown by red in **(I-K)** and fire in **(I’-K’)**, where white indicates high ROS (1) and black-blue indicates low ROS levels (0) in lymph gland progenitor cells**. (I, I’)** control (RF, *domeMeso-Gal4,UAS-GFP/+*) lymph gland showing ROS levels in lymph gland progenitor-cells (area demarcated within the yellow border). While, expressing **(J, J’)** *Gat^RNAi^* (RF, *domeMeso-Gal4,UAS-GFP;UAS-Gat^RNAi^*) leads to increase in progenitor ROS as compared to control **(I, I’)**, supplementing this genetic condition with **(K, K’)** serine (Serine, *domeMeso-Gal4,UAS-GFP;UAS-Gat^RNAi^*) recovers the increased ROS levels. For comparison, also refer to **(J, J’)** *Gat^RNAi^* raised on regular food (RF). For quantifications, refer to **N**. **(L)** Quantification of blood-progenitor cysteine (Cys) levels (fold change, f.c.) in *domeMeso>GFP/+* (control, RF, n=16)*, domeMeso>GFP/Gat^RNAi^* (RF, n=13, p=0.0132), and *domeMeso>GFP/Gat^RNAi^* (Serine, n=18, p=0.0754 in comparison to control and p=0.0017 in comparison to *Gat^RNAi^*, RF). **(M)** Quantification of blood-progenitor GSH levels (fold change, f.c.) in *domeMeso>GFP/+* (control, RF, n=17)*, domeMeso>GFP/Gat^RNAi^* (RF, n=14, p=0.0003), and *domeMeso>GFP/Gat^RNAi^* (Serine, n=16, p=0.0942 in comparison to control and p=0.0308 in comparison to *Gat^RNAi^*, RF). **(N)** Quantification of blood-progenitor ROS levels (fold change, f.c.) in *domeMeso>GFP/+* (control, RF, n=25)*, domeMeso>GFP/Gat^RNAi^* (RF, n=16, p=0.0088), and *domeMeso>GFP/Gat^RNAi^* (Serine, n=11, p=0.0161 in comparison to control and p=0.6800 in comparison to *Gat^RNAi^*, RF). **(O)** Quantification of lymph gland area in *domeMeso>GFP/+* (control, RF, n=25)*, domeMeso>GFP/Gat^RNAi^* (RF, n=17, p<0.0001), and *domeMeso>GFP/Gat^RNAi^* (Serine, n=15, p=0.0127 in comparison to control and p=0.0158 in comparison to *Gat^RNAi^*, RF).

In this context, intracellular availability of serine and any correlation with pyruvate metabolism was explored. Serine levels can be managed either via its uptake through transporters (47) or through *de novo* routes that generate it (48). In this regard, *de novo* synthesis becomes interesting, because pyruvate via gluconeogenesis can be a predominant source to generate serine (49). It is therefore possible that increased pyruvate oxidation in *Gat* mutants impaired pyruvate availability for gluconeogenesis and consequently serine biosynthesis. To understand this, we undertook a U^13^C-pyruvate isotope based metabolic labelling analysis in wandering 3^rd^ instar control and GatRNAi lymph glands and assessed pyruvate metabolism.

Pyruvate is a 3 carbon acid and contribute its two carbons to the TCA cycle through the pyruvate dehydrogenase (PDH) dependent generation of acetyl-CoA, which combines with oxaloacetate to form citrate and other TCA metabolites (Fig. 4A). Pyruvate can additionally fuel its full 3 carbons to generate oxaloacetate through the activity of pyruvate carboxylase (PC) (50), which lies at the centre of gluconeogenesis (51,52). Thus, the flow of labels between the two routes would indicate how lymph gland cells utilized pyruvate. The m+2 label in oxaloacetate is derived from PDH, while m+3 label incorporation is mediated by PC. As various steps of TCA cycle are reversible and also feed cyclically, 3 carbon label (m+3) contributed from PC derived OAA conversion and m+2 label contributed from PDH derived acetyl-CoA can lead to higher ^13^C incorporation as the cycle continues and is indicative about the overall TCA activity (Fig. 4A).

**Figure 4.**
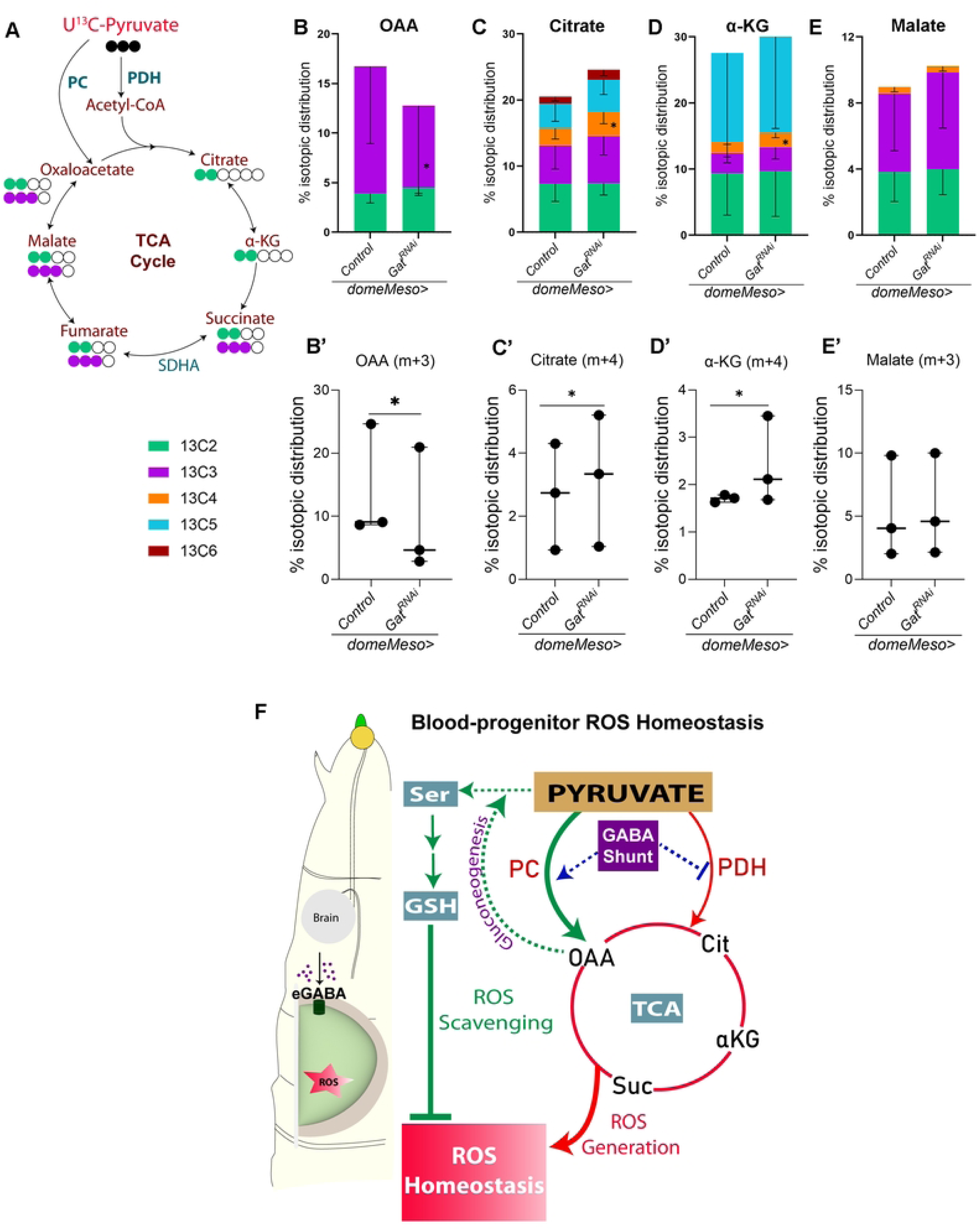
GABA catabolism regulates pyruvate metabolism to maintain blood-progenitor ROS homeostasis. Data is presented as mean with SD and stacked bar plots for **B-E** and median plots (*p<0.05) for **B’-E’**. Two-way ANOVA is applied for **B’-D’**. In **B’-E’,** ‘N’ is number of experimental repeats and ‘n’ is sample size and shown by black dot (N). U^13^C-universal 13C, all three carbons are ^13^C, PDH-pyruvate dehydrogenase, PC-pyruvate carboxylase, SDHA-Succinate dehydrogenase A, Ser-serine, GSH-glutathione, Cit-citrate, α-KG-α-ketoglutarate, Suc-succinate, OAA-oxaloacetate, GABA-γ-aminobutyric acid, eGABA-extracellular GABA, TCA-tricarboxylic acid, ROS-reactive oxygen species. **(A)** Schematic representation of the U^13^C-pyruvate (black) isotopic labelling. U_13_C-pyruvate contributes two 13C carbons (green) to TCA metabolites via PDH and three carbons (purple) via PC mediated metabolism to OAA, which further cycles and contributes to TCA cycle metabolites. **(B-E’)** Percentage (%) isotopic distribution of U^13^C-pyruvate into TCA cycle metabolites, where isotopic distribution is depicted in different colours and m+2 is green, m+3 is purple, m+4 is orange, m+5 is blue and m+6 is dark red. **(B)** Control (*domeMeso>GFP/+*, n=8) and *domeMeso>GFP/Gat^RNAi^* (n=8) lymph glands showing percentage label incorporation for m+2 and m+3 in OAA, **(B’)** lymph glands expressing *Gat^RNAi^* in the progenitor cells (*domeMeso>GFP/Gat^RNAi^*, N=3, n=8, p=0.0455) show a significant reduction for m+3 percentage label incorporation in OAA as compared to control (*domeMeso>GFP/+*, N=3, n=8), **(C)** control (*domeMeso>GFP/+*, n=8) and *domeMeso>GFP/Gat^RNAi^* (n=9) lymph glands showing percentage label incorporation for m+2 to m+6 (higher isotopes) in citrate, **(C’)** lymph glands expressing *Gat^RNAi^* in the progenitor cells (*domeMeso>GFP/Gat^RNAi^*, N=3, n=9, p=0.0456) show a significant increase for m+4 percentage label incorporation in citrate as compared to control (*domeMeso>GFP/+*, N=3, n=8), **(D)** control (*domeMeso>GFP/+*, n=8) and *domeMeso>GFP/Gat^RNAi^* (n=9) lymph glands showing percentage label incorporation for m+2 to m+5 (higher isotopes) in α-KG, **(D’)** lymph glands expressing *Gat^RNAi^* in the progenitor cells (*domeMeso>GFP/Gat^RNAi^*, N=3, n=9, p=0.0262) show a significant increase for m+4 percentage label incorporation in α-KG as compared to control (*domeMeso>GFP/+*, N=3, n=8) and **(E)** control (*domeMeso>GFP/+*, n=8) and *domeMeso>GFP/Gat^RNAi^* (n=9) lymph glands showing percentage label incorporation for m+2 to m+4 (higher isotopes) in malate, **(E’)** lymph glands expressing *Gat^RNAi^* in the progenitor cells (*domeMeso>GFP/Gat^RNAi^*, N=3, n=9, p=0.5419) does not show any change for m+3 percentage label incorporation in malate as compared to control (*domeMeso>GFP/+*, N=3, n=8). **(F) Control of *de novo* GSH synthesis in lymph gland blood progenitor cells by the GABA-shunt pathway.** Pyruvate is metabolised into OAA by PC and PDH. GABA controls PDH activity and restricts TCA cycle. This state is necessary to support GSH production in the progenitor cells. We speculate pyruvate’s availability for gluconeogenesis, as the key step to allow *de novo* synthesis of serine. Serine is a key intermediate for promoting GSH production in these cells. GABA by positively influencing serine production and regulating PDH activity, enables a state in blood progenitors cells, whereby they are able to generate developmentally regulated ROS and also capacitate them with ROS scavenging potential and counter any redox stress. This metabolic program maintains homeostatic ROS levels in the lymph gland, where ROS sensitizes progenitors to differentiation cues (5), but also limits its excessive production to support lymph gland growth (25).

In control lymph glands, 15% total labelling was seen in OAA. Of this, approximately 10% was m+3 ^13^C labelled, which is PC-derived and 5% was m+2, which is PDH-derived (Fig. 4B). This showed PC as a favoured route for pyruvate utilization over PDH in control conditions. When assessed for TCA metabolites, we observed m+2 derived ^13^C-pruvate label in citrate, α-KG and malate (Fig. 4C-E). Moreover, we also detected higher order label of ^13^C in these metabolites (Fig. 4C-E) and the dynamics demonstrated pyruvate’s contribution to fueling the TCA cycle. The data revealed a functional TCA, but regulated and aligned with our genetic findings that showcased GABA dependent regulation of PDH activity.

We conducted the same analysis in *domeMeso>Gat^RNAi^* lymph glands where the U^13^C-pyruvate label incorporation analysis revealed a rather unexpected reduction in PC derived m+3 OAA label incorporation (Fig. 4B, B’). PDH derived m+2 OAA label, however, remained comparable to the control (Fig. 4B). Importantly, we observed increase in overall label incorporation in TCA metabolites with higher m+4 isotopic carbons detected for citrate and α-KG (Fig. 4C-D’). The pyruvate flux towards malate in *Gat^RNAi^* condition however remained comparable to control (Fig. 4E-E’). The significant reduction in labelled OAA, but the concomitant increase in higher label incorporation in TCA metabolites in *domeMeso>Gat^RNAi^* lymph glands implied heightened flux of fueling intermediates into the TCA cycle. This metabolic data corroborated with our genetic findings (25) and reinstated the importance of GABA breakdown in controlling overall TCA activity. The data also revealed an unexpected preference for PC as opposed to PDH in the control lymph glands and the influence of GABA metabolism in this context. This allows us to speculate that in homeostasis, the m+3 labelled OAA derived from PC activity, through gluconeogenic routes drives serine generation in progenitor cells. This route to *de novo* synthesize serine most likely capacitates the progenitor cells with better control on their redox state. In the absence of GABA metabolism, the reduction in the flux of PC, with the concomitant elevation in PDH activity, leading to elevated TCA rate, the majority of OAA is incorporated into generating citrate. As a result, this state leads to impairment of serine generation and subsequently GSH synthesis. The multifaceted control exerted by GABA shunt on pyruvate’s metabolic, where it positively influences PC and LDH while negatively influences PDH seems like a metabolic framework that is established by GABA in the progenitor cells that keeps their overall development and redox homeostasis in place.

## Discussion

Blood progenitor cells in the lymph gland rely on reactive oxygen species (ROS) to maintain their differentiation potential, but they must also regulate redox balance to sustain an undifferentiated state and avoid precocious differentiation (5). This interplay between metabolic processes that facilitate ROS homeostasis forms the central focus of this study.

### Proposed Model

Based on our findings, including mass spectrometry data and genetic studies, we propose the following model. In the lymph gland, pyruvate is metabolized into oxaloacetate (OAA) primarily via pyruvate carboxylase (PC) and to a lesser extent through pyruvate dehydrogenase (PDH). PC is favoured over active PDH, tightly regulating OAA pools and the rate of TCA cycling. This balance enables the system to generate developmentally regulated ROS while preserving OAA for additional processes, such as the gluconeogenic production of serine, cysteine, and glutathione (GSH). These metabolites, in turn, contribute to the antioxidant capacity of blood progenitor cells.

GABA catabolism regulates PDH activity and influences PC function, facilitating serine synthesis and thereby maintaining a finely tuned redox state. In the absence of GABA, pyruvate is preferentially utilized via PDH, leading to excessive ROS production and serine deprivation. Reducing PDH activity restores pyruvate dynamics, reestablishing cysteine and GSH levels. Overall, this highlights an essential nexus in blood progenitor cells, where ROS generation is integrally linked to ROS scavenging, with GABA metabolism centrally regulating pyruvate’s availability at this junction (Fig. 4F).

### TCA Cycle, GABA-Shunt and Redox Regulation

The TCA cycle is a critical metabolic hub that modulates ROS levels in progenitor cells. Pyruvate’s entry into the TCA cycle through PDH, a rate-limiting enzyme that converts pyruvate to acetyl-CoA, is the primary route for ROS production in these cells (25). Blood progenitor cells tightly regulate PDH activity to prevent excessive ROS accumulation. Previous studies have demonstrated that GABA catabolism plays a key role in this regulation, with succinate derived from GABA metabolism modulating PDH activity and maintaining ROS homeostasis.

Our metabolic flux measurements reveal PC as an additional route for pyruvate metabolism, which is impaired in the absence of GABA. GABA catabolism fine-tunes TCA cycling by regulating PDH activity and limiting intermediate utilization, such as OAA, within the TCA. OAA’s availability is necessary for gluconeogenic programs, including serine biosynthesis. In GABA mutants, increased PDH activity prioritizes intermediate utilization, leading to a lack of OAA for other metabolic processes, creating a bottleneck that compromises antioxidant synthesis. Accelerated TCA cycling depletes intermediates needed for serine and GSH biosynthesis, tipping the redox balance toward oxidative stress.

Dysregulation of TCA flux, such as succinate accumulation, has been linked to impaired redox homeostasis (53). The GABA shunt, in this regard, is often seen as a bypass of the conventional TCA cycle, where it plays a significant role in TCA regulation. GABA is metabolized into succinate through the sequential action of GABA transaminase and succinate semialdehyde dehydrogenase (SSADH). Succinate enters the TCA cycle directly, bypassing the key steps like α-KDH and PDH that contribute to ROS generation. Thus, by providing an alternate source of TCA intermediates, the GABA shunt fine-tunes the rate of TCA cycling and mitigates excessive ROS production. However, in the lymph gland blood progenitor cells the catabolism of GABA into succinate functions in a completely different way. We find GABA-derived succinate to be closely associated with its role in the post-translational regulation of protein function. Either through regulation of Hph and thereby stabilizing Hif-1α (26), or by controlling PDK activity and PDH phosphorylation (25), GABA shunt operates to control key enzymatic steps that are rate limiting to either glycolysis or TCA activity. The influence of shunt in the production of OAA by regulating PC activity, is yet another revelation in this study and positions GABA shunt pathway at the nexus of ensuring a balance between energy production and biosynthetic needs within the progenitor cells.

### Blood progenitor reliance on Serine/GSH de novo biosynthesis: a means to enable adaptability to stress conditions

Serine, a precursor for GSH synthesis, is produced from glycolytic intermediates and is directly influenced by TCA activity. TCA-derived α-ketoglutarate and malate promotes serine synthesis by activating key enzymes in the pathway, such as phosphoglycerate dehydrogenase (PHGDH) (54). PHGDH activity is sensitive to the NAD/NADH ratio, which TCA metabolites maintain through pyruvate-to-lactate conversion and the malate-aspartate shuttle (55). In macrophages, dysregulation of *de novo* serine synthesis, impairs GSH synthesis and ROS homeostasis leading to loss of inflammatory response in them (56).

*De novo* serine synthesis in blood progenitor cells and its contribution to GSH synthesis by serving as a carbon donor for cysteine synthesis is very striking. Moreover, the influence of GABA sustaining this state is even more impressive. We know that in the blood progenitor cells, GABA catabolism through stabilizing Hif-1α and LDH activity, promotes their glycolytic state, which is necessary to support their immune competency (26,57). Thus GABA functions in immune modulation and maintains a pro-inflammatory state in progenitor cells. While the sustenance of a glycolytic state may allow the NAD/NADH ratios to be maintained to support serine synthesis, the implications of maintaining a *de novo* route to GSH generation may be a part of the larger immune competency program that GABA catabolism supports. The purpose of utilizing glycolytic intermediates to make serine and GSH, may be an option to counter oxidative stresses during infection and avoid any progenitor cell death. Serine metabolism is also closely linked to one-carbon metabolism where it integrates nutritional status and generates diverse outputs, including methylation substrates (58). For example, glucose-dependent serine biosynthesis supports T-cell proliferation (59), while TCA-dependent serine synthesis via PEPCK2 is central to metabolic adaptation in tumor cells (60). Thus keeping this route to generate serine and GSH may also allow the progenitor cells to maintain a metabolic state that integrates inputs from multiple nutritional routes, ensuring adaptability and resilience under oxidative environmental stressors.

### Implications for Hematopoietic Systems and conclusions

These blood progenitor cells of the lymph gland, like the common myeloid precursors are highly dependent on stress signals for their development. They develop in niches with high ROS (5,21), high calcium (13,61) and respond promptly to any changes in their levels. Thus, these cells have optimised the use of stress signals to program their development. In this regard, our study alludes to the underlying progenitor-specific metabolic program that ensures their adaptability and resilience to such developmental stressors. This notion is supported by multiple pieces of published evidences that show how impairing GABA function in progenitor cells, affects their ability to respond to environmental changes, that include sensory stimulation, infection and also changes to their local niche environment, like alteration in calcium (13) or ROS homeostasis (25). The control of GABA shunt in optimising pyruvate metabolism, to ensure a multifaceted development of the blood-progenitor cells is indeed the most prominent phenomenon that emerges from our past and current study.

Mammalian hematopoietic progenitor cells exhibit a similar reliance on serine biosynthesis (62) and GABA/GABA-receptor signaling (34). ROS signaling plays a crucial role in hematopoietic stem cell (HSC) proliferation, with NAC treatments restoring HSC quiescence and proliferation under oxidative stress (35). These commonalities lead us to speculate similar modalities for metabolic regulation in mammalian hematopoietic stem cell niches as well. The optimal application of this metabolic framework to safeguard differentiation potential under oxidative stress is striking and could be a conserved phenomenon that however remains to be investigated. Overall, the intricate interplay between metabolic and redox regulation during development and the integral role of GABA shunt in keeping the checks in place positions GABA in an entirely new developmental context, as a redox modulator in stem/progenitor cells.

## Acknowledgements

We thank the Bloomington *Drosophila* Stock Center (BDSC) and Vienna *Drosophila* Resource Center (VDRC) for fly stocks. We acknowledge the support of Central Imaging & Flow Cytometry Facility (CIFF) and the fly facility at National Centre for Biological Sciences (NCBS), Centre for Cellular and Molecular Platforms (C-CAMP). We thank metabolomics facility at MPF, Bindley Bioscience Center for help with metabolomics experiments and Dr. Vikki Weake, Purdue University for providing the space for *Drosophila* related work. We thank Patrick Jouandin and Norbert Perrimon for their suggestions and feedback on metabolomics experiments. Owing to space limitations, we apologize to our colleagues whose work is not cited. This study was supported by Department of Science and Technology - CRG (Grant number CRG/2021/002815), USIAS Indo French grant awarded to T.M and DBT/Wellcome Trust Senior Research Fellowship (Grant number IA/S/22/1/506259) awarded to T.M. M.G. is a graduate student at inStem, in the Mukherjee lab, and is supported by DST-INSPIRE and, SERB-OVDF fellowship facilitated the work done at Purdue University. S.T. is a graduate student at inStem, in the Mukherjee lab, and is supported by UGC-JRF.

## Materials and Methods

### Drosophila husbandry, stocks, and genetics

The following *Drosophila melanogaster* stocks were used in this study: *w^1118^* (wild type*, wt*) and *domeMeso-Gal4,UAS-GFP* (U. Banerjee). The *RNAi* stocks were either obtained from VDRC (Vienna) or BDSC (Bloomington) *Drosophila* stock centres. The lines used in this study are: *Gat^RNAi^* (BDSC 29422)*, Ssadh^RNAi^* (VDRC 106637)*, Pdha^RNAi^* (BDSC 55345) and *Pdk^RNAi^* (BDSC 28635).

All fly stocks were reared on corn meal agar food medium with yeast supplementation at 25°C incubator unless specified. Tight collections were done for 4-6 hours to avoid over-crowding and for synchronous development of larvae. The crosses involving RNA*i* lines were maintained at 29°C to maximize the efficacy of the *Gal4/UAS* RNA*i* system. Controls correspond to *Gal4* drivers crossed with *w^1118^*.

### Immunostaining and immunohistochemistry

Immunohistochemistry on lymph gland tissues were performed with the following primary antibodies: rabbit-αGlutathione (GSH, 1:100, abcam #ab9443), mouse-αGlutamate (Glu, 1:100, Sigma G9282) and mouse-αCysteine (Cys, 1:20, sc-69954). The secondary antibodies Alexa Flour 488, 546 and 647 (Invitrogen) were used at 1:400 dilutions. Nuclei were visualized using DAPI (Sigma). Samples were mounted with Vectashield (Vector Laboratories).

Lymph glands dissected from wandering 3^rd^ instar larvae were stained following the protocol of (63). Lymph gland tissues from synchronized larvae of required developmental stage were dissected in cold PBS (1X Phosphate Buffer Saline, pH-7.2) and fixed in 4% formaldehyde for 40 min. at room temperature. Tissues were then washed thrice (15 min. each wash) in 0.3% PBT (0.3% triton-X in 1X PBS) for permeabilization and further blocking was done in 5% NGS, for 45 min at RT. After this, tissues were incubated in the respective primary antibodies with appropriate dilution in 5% NGS overnight at 4°C. Post primary antibody incubation, tissues were washed thrice in 0.3% PBT for 15 min each. This was followed by incubation of tissues in respective secondary antibodies for 2-3 hrs at RT. After secondary antibody incubation, tissues were washed in 0.3% PBT for 15 min. following a DAPI+0.3% PBT wash for 15 min. Excess DAPI was washed off by a wash of 0.3% PBT for 15 min. Tissues were mounted in Vectashield (Vector Laboratories) and then imaged utilizing confocal microscopy. For representation, one lymph gland lobe is shown in the figure panels.

### ROS (DHE) detection in lymph glands

Lymph glands dissected from the wandering 3^rd^ instar larvae were stained for ROS levels following the protocol of (64). The dissected lymph gland tissues were stained in 1:1000 DHE (Invitrogen, Molecular Probes, D11347) dissolved in 1X PBS for 15 min in the dark. Tissues were washed in 1X PBS twice and fixed with 4% formaldehyde for 6-8 min. at room temperature in the dark. Tissues were again quickly washed in 1X PBS twice and then mounted in Vectashield (Vector Laboratories). The lymph glands were imaged immediately.

### Image acquisition and processing

Immuno-stained and DHE stained (ROS) lymph gland tissue images were acquired using Olympus FV3000 Confocal Microscopy 40X oil-immersion objective. Microscope settings were kept constant for each sample in every experiment. The image acquisition settings were chosen to capture this difference in control lymph glands without causing saturation in majority of the pixels. This setting was thereafter kept constant for all other genotypes that were conducted in the corresponding experimental batch and were processed for analysis and quantifications. Lymph gland images were processed using ImageJ (Fiji) and Adobe Photoshop software.

### Quantification of lymph gland phenotypes

All images were quantified using ImageJ (Fiji) software and Microsoft Excel. Images were acquired as z-stacks and quantifications were done as described previously (25). For antibody intensity and ROS levels quantifications, the area covering the dome^+^ region of the lymph gland lobe was marked utilizing free hand tool in ImageJ and mean fluorescence intensity was calculated. Background noise was quantified marking four random square of equal size in each image and corresponding average intensities were subtracted from the mean intensity values calculated from the respective signal. The relative fold change was calculated from the final mean fluorescence intensity values in Microsoft excel and graph plotting, and statistical data analysis was performed using GraphPad Prism software. For all intensity quantifications, the laser settings for each individual experimental set-up were kept constant and controls were analysed in parallel to the mutant conditions every time. For lymph gland area analysis, middle two z-stacks of the image were merged, and total lymph gland lobe area was marked using the free-hand tool of ImageJ and then analysed further for quantifications. In all experiments “n” implies the total number of lymph gland lobes analysed which were obtained from multiple independent experiments, repeated at least three times.

### Metabolite supplementation

Glutathione reduced (GSH, G4251), Succinate (SF, Sodium succinate dibasic hexahydrate, Sigma, S9637), N-Acetyl-L-cysteine (NAC, Sigma, A7250), serine (Sigma, S4500) and methionine (Met, Sigma, M9625) enriched diets were prepared by supplementing regular fly food with weight/volume measures of each supplement and 0.1% GSH, 3% Succinate, 0.1% NAC, 0.1% methionine and 0.1% serine concentrations were used. Eggs were transferred in these supplemented diets and reared until analysis of the lymph gland tissues at the wandering 3^rd^ instar stage.

### TCA metabolite extraction and targeted metabolomic analysis

For metabolite extraction, five lymph glands from wandering 3^rd^ instar larvae were taken per sample in 200 μl of 80% ice-cold methanol and samples were homogenized briefly. Samples were stored at −80 degrees immediately and later dried down in a Vacufuge plus speed-vac at room temperature and derivatized further with OBHA/EDC for metabolite analysis (65,66). For derivatization, dried samples were dissolved in 50 μl of LC/MS grade water and 50 μl of 1M EDC (in Pyridine buffer, pH 5) was added. These samples were kept on a thermomixer for 10 min. at room temperature and 100 μl of 0.5M OBHA (in Pyridine buffer, pH 5) was added. The samples were incubated again for 1.5 hours on the thermomixer at 25°C, and metabolites were extracted by adding 300 μl of ethyl acetate and this step was repeated three times. Samples were dried down in a Vacufuge plus speed-vac at room temperature and stored at −80°C until run for LC/MS analysis. The metabolite extract was separated using a Waters XBridge C18 Column (2.1 mm, 100 mm, 3.5 mm) on an Agilent QQQ 6470 system coupled to a Agilent 1290 UPLC system. The autosampler and column oven were kept at 4°C and 25°C, respectively. The buffers utilised for the analysis were, buffer A: Water plus 0.1% Formic Acid and buffer B 100% acetonitrile (ACN) plus 0.1% Formic Acid. A flow rate of 0.300 ml/minute was used for the chromatographic gradient as follows: 0 min-10% B; 0.50 min: 10% B; 8 min: 100% B; 10 min: 10% B; 11 min: 10% B and at 16 min gradient was held at 10%B. MRM, positive ion mode was used for running the LC/MS and mass spectrometry detection was carried out on a QQQ Agilent 6470 system with ESI source attached to a UPLC system. For metabolite quantification, Peak areas were processed using MassHunter workstation (Agilent). Microsoft Excel and GraphPad Prism software were used for statistical analysis.

### Amino acid extraction and LC/MS/MS analysis

For amino acid (serine) analysis, five lymph glands from wandering 3^rd^ instar larvae were taken and TCA (trichloroacetic acid) precipitation method was utilized. Briefly, lymph glands were dissected in 200μl of 1X PBS (Gibco) and 40μl of TCA (100% TCA soln.) was added. Samples were homogenized and incubated in ice for 10 minutes. After this, samples were centrifuged at 13000 RPM for 10 minutes at 4 degrees. Supernatant was transferred to fresh Eppendorf tubes and stored at −80 degrees until analysis. LC/MS/MS based analysis for amino acids was done utilizing HILIC column on Agilent UPLC-QQQ3 6470 system.

### 13C-isotopic labelling and stable isotope tracer analysis for metabolite measurements

For isotopomer tracer analysis, wandering 3^rd^ instar larvae were washed twice in PBS and lymph glands were dissected. Larval lymph glands were incubated in 10mM of U^13^C-Pyruvate (Cambridge Isotope Laboratories, CLM-2440) in 1X PBS (Gibco) for 30 min. at room temperature. Samples were quickly rinsed in LC/MS grade water and processed for metabolite extraction and derivatization for TCA metabolite analysis.

For LC/MS based steady-state and flux analysis five lymph glands per replicate (n) were taken and each experiment was repeated a minimum of three times (N).

### Statistical analyses

All statistical analyses and quantifications were performed using GraphPad Prism 10 and Microsoft Excel 2016. Mann-Whitney test is employed for statistical significance for all the experiments except the LC/MS based analysis, where two-way ANOVA is applied to determine significance. *Drosophila* are not limiting, therefore no power calculations were used to pre-determine sample size. In box and whisker plots, the horizontal line indicates the median, whiskers indicate the minimum and maximum values, and the box indicates the lower and upper quartiles. Statistical analyses and sample size were approached with the same level of rigor as done in our previous study (25). Plots, test applied and sample size is mentioned separately in the figure legends section.

## Supplementary Figure Legends

## Supplementary Figure

**Figure S1.**
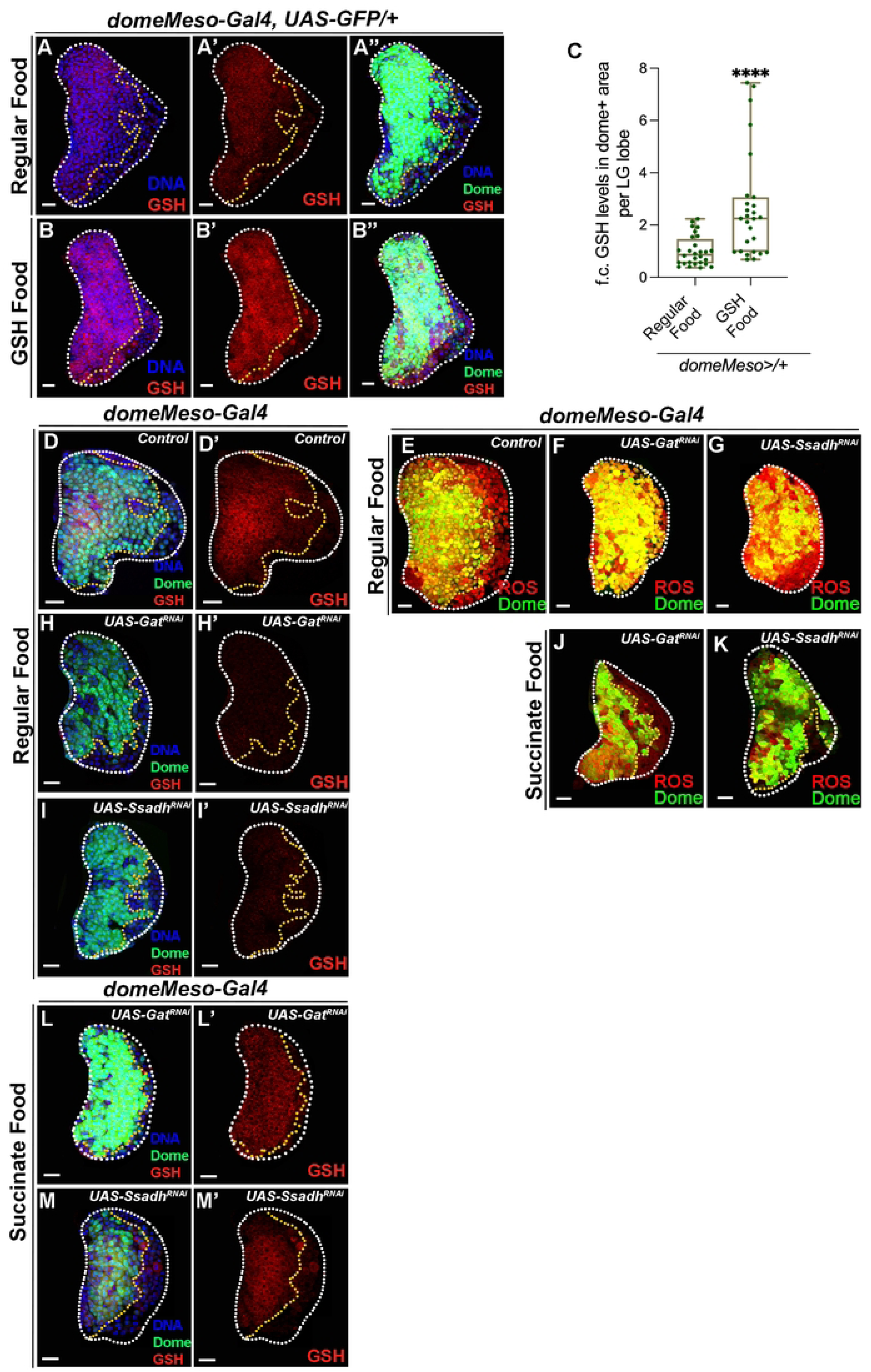
GABA catabolic pathway in *Drosophila* blood-progenitor cells control their GSH levels. RF is regular food, and GSH and succinate food is GSH and succinate supplemented food respectively. Data is presented as median plots (****p<0.0001). Mann-Whitney test is applied for **C**. In **C**, ‘n’ is total number of lymph gland lobes analysed and in the graph each lobe is represented by a green dot. DNA is stained with DAPI in blue, dome marks the progenitor cells in green. Comparisons for significance are done with their respective control. White border demarcates the lymph gland lobe and yellow border marks the dome positive area towards the left side. **(A-B”)** Representative images showing GSH levels in the lymph gland progenitor cells shown by merge of dome+ (green), DNA (blue) and GSH (red) from *domeMeso-Gal4,UAS-GFP/+* genetic backgrounds. Control (RF, *domeMeso-Gal4,UAS-GFP/+*) showing **(A-A”)** GSH (red) levels in the progenitor cells and supplementing this genetic condition with GSH (GSH, *domeMeso-Gal4,UAS-GFP/+*) leads to a significant increase in GSH levels in the progenitor cells. For comparison, refer to control (Regular Food) **A-A”**. For quantifications, refer to **C**. **(C)** Quantification of blood-progenitor GSH levels in *domeMeso>GFP/+* (control, Regular Food, n=28)*, and domeMeso>GFP/+* (GSH Food, n=27, p<0.0001). **(D-I’)** Representative images showing ROS and GSH levels in the lymph gland progenitor cells shown by merge of dome+ (green), DNA (blue) and respective expression (GSH/ROS, red) from different genetic backgrounds. Control (*domeMeso-Gal4,UAS-GFP/+*) showing elevated **(D, D’)** GSH (red) levels and **(E)** ROS (stained with DHE, red) levels in the lymph gland progenitor cells, expressing **(F)** *Gat^RNAi^* (RF, *domeMeso-Gal4,UAS-GFP;UAS-Gat^RNAi^*) and **(G)** *Ssadh^RNAi^* (RF, *domeMeso-Gal4,UAS-GFP;UAS-Ssadh^RNAi^*) leads to an increase in blood-progenitor ROS levels as compared to control **(B)** and, expressing **(H, H’)** *Gat^RNAi^* (RF, *domeMeso-Gal4,UAS-GFP;UAS-Gat^RNAi^*), **(I, I’)** *Ssadh^RNAi^* (RF, *domeMeso-Gal4,UAS-GFP;UAS-Ssadh^RNAi^*) leads to a decrease in blood-progenitor GSH levels as compared to control **(D, D’)**. **(J-M’)** Succinate supplementation to **(J, L, L’)** *Gat^RNAi^* (SF, *domeMeso-Gal4,UAS-GFP;UAS-Gat^RNAi^*) and **(K, M, M’)** *Ssadh^RNAi^* (SF, *domeMeso-Gal4,UAS-GFP;UAS-Ssadh^RNAi^*) leads to rescue of blood-progenitor **(J, K)** ROS and **(L-M’)** GSH defect. For comparison, refer to *Gat^RNAi^* **(F, H, H’)** and *Ssadh^RNAi^* **(G, I, I’)** raised on regular food.

**Figure S2.**
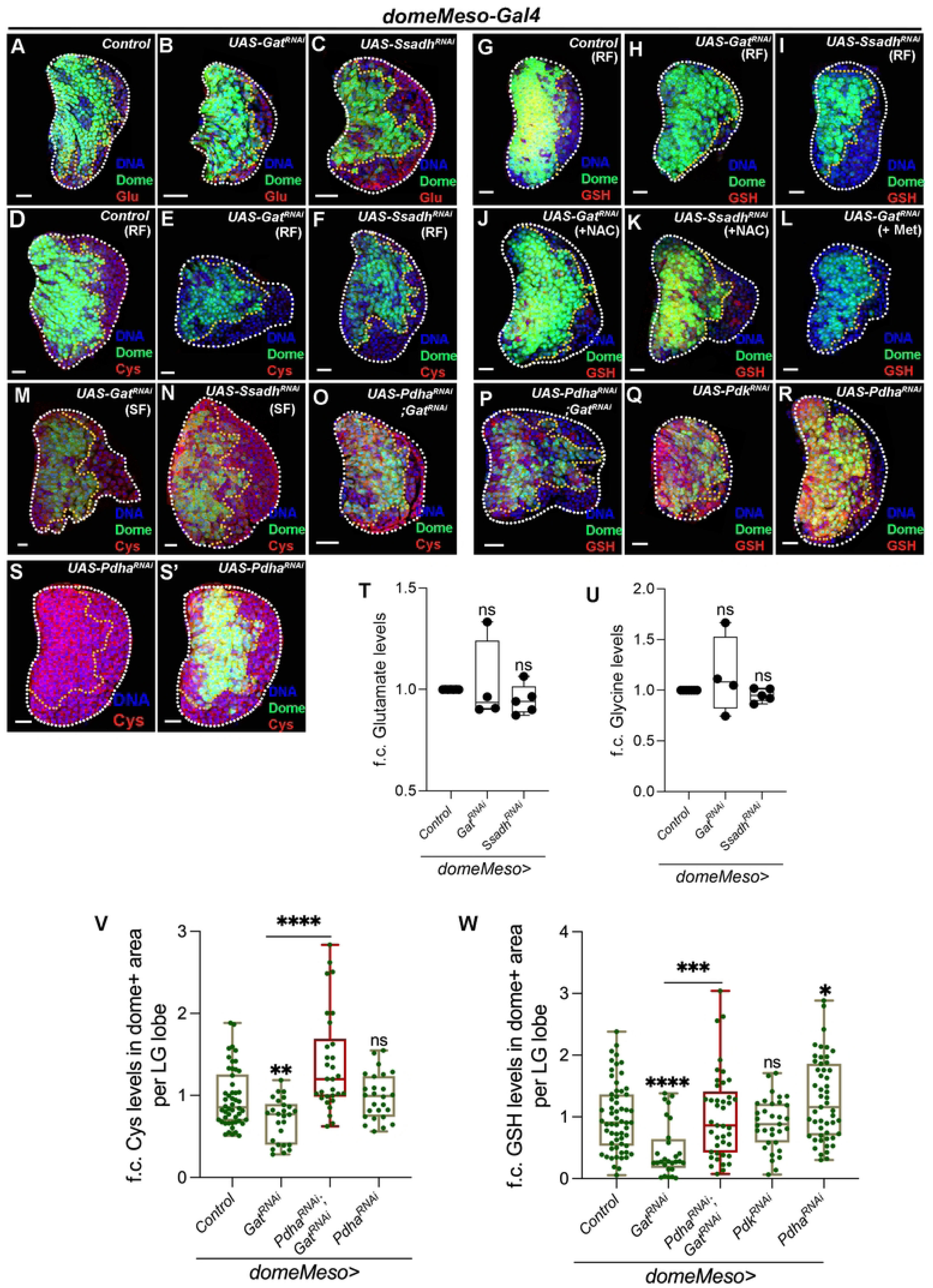
Regulation of glutathione synthesis by GABA catabolic pathway. RF is regular food, SF is succinate food and NAC is N-acetylcysteine supplemented food. DAPI marks DNA. Dome (green) marks progenitor cells. Data is presented as median plots (*p<0.05; **p<0.01; ***p<0.001; ****p<0.0001 and ns is non-significant). Two-way ANOVA, Dunnett’s multiple comparison test is applied for **T, U** and Mann-Whitney test is applied for **V, W**. In **T, U** ‘N’ is number of experimental repeats and ‘n’ is sample size and shown by black dot (N). In **V, W** ‘n’ is total number of lymph gland lobes analysed and in the graph each lobe is represented by a green dot. Scale bar: 20µm. DNA is stained with DAPI in blue, dome marks the progenitor cells in green. Comparisons for significance are done with their respective control and also with respective genetic conditions which are indicated by horizontal lines drawn above the boxplots. Red bars represent rescue combinations. White border demarcates the lymph gland lobe and yellow border marks the dome positive area towards the left side. **(A-C)** Representative images showing glutamate expression in lymph gland progenitor cells (area marked within the yellow dotted line) with merge of dome+ (green), DNA (blue) and glutamate (Glut, red) across different genetic backgrounds. In comparison to **(A)** control (*domeMeso-Gal4,UAS-GFP/+*) lymph gland showing uniform glutamate levels across all cells of the tissue, including the progenitor-cells (area within the yellow dotted line). Expressing **(B)** *Gat^RNAi^* (*domeMeso-Gal4,UAS-GFP;UAS-Gat^RNAi^*) and **(C)** *Ssadh^RNAi^* (*domeMeso-Gal4,UAS-GFP;UAS-Ssadh^RNAi^*) in the progenitor cells does not affect their glutamate levels. **(D-F)** Representative images showing cysteine (Cys, red) expression in lymph gland progenitor cells (area marked within the yellow dotted line) with merge of dome+ (green), DNA (blue) and cysteine (red) from different genetic backgrounds. **(D)** Control (RF, *domeMeso-Gal4,UAS-GFP/+*) lymph gland showing relatively uniform cysteine levels in all cells of the lymph gland including progenitor-cells (area demarcated within the yellow border). Expressing **(E)** *Gat^RNAi^* (RF, *domeMeso-Gal4,UAS-GFP;UAS-Gat^RNAi^*) and **(F)** *Ssadh^RNAi^* (RF, *domeMeso-Gal4,UAS-GFP;UAS-Ssadh^RNAi^*) in the progenitor cells leads to reduction in cysteine levels as compared to control **(D)**. **(G-L)** Representative images showing GSH (red) expression in lymph gland progenitor cells (area marked within the yellow dotted line) with merge of dome+ (green), DNA (blue) and GSH (red)from different genetic backgrounds. **(G)** Control (RF, *domeMeso-Gal4,UAS-GFP/+*) lymph gland showing GSH levels. While expressing **(H)** *Gat^RNAi^* (RF, *domeMeso-Gal4,UAS-GFP;UAS-Gat^RNAi^*) and **(I)** *Ssadh^RNAi^* (RF, *domeMeso-Gal4,UAS-GFP;UAS-Ssadh^RNAi^*) in the progenitor cells leads to reduction in GSH levels, supplementing these genetic conditions with **(J, K)** N-acetyl cysteine (NAC), restores GSH levels in both **(J)** *Gat^RNAi^* (NAC, *domeMeso-Gal4,UAS-GFP;UAS-Gat^RNAi^*) and **(K)** *Ssadh^RNAi^* (NAC, *domeMeso-Gal4,UAS-GFP;UAS-Ssadh^RNAi^*) background to levels almost comparable to control shown in **G**. **(L)** Methionine supplementation to *Gat^RNAi^* (Met, *domeMeso-Gal4,UAS-GFP;UAS-Gat^RNAi^*) does not recover GSH levels. For comparison also refer to *Gat^RNAi^* **(J)** and *Ssadh^RNAi^* **(K)** raised on regular food (RF). **(M-N)** Succinate supplementation to **(M)** *Gat^RNAi^* (SF, *domeMeso-Gal4,UAS-GFP;UAS-Gat^RNAi^*) and **(N)** *Ssadh^RNAi^* (SF, *domeMeso-Gal4,UAS-GFP;UAS-Ssadh^RNAi^*) restores blood-progenitor cysteine levels. For comparison see lymph gland lobes from **(E)** *Gat^RNAi^* and **(F)** *Ssadh^RNAi^* animals raised on regular food (RF) and control in **D**. **(O-P)** Progenitor specific expression of *Pdha^RNAi^* in *Gat^RNAi^* animals (*domeMeso-Gal4,UAS-GFP; UAS-Pdha^RNAi^;UAS-Gat^RNAi^*) leads to a recovery of blood-progenitor **(O)** cysteine and **(P)** GSH levels. Compare with **(E)** Cys and **(H)** GSH in *Gat^RNAi^ (domeMeso-Gal4,UAS-GFP;UAS-Gat^RNAi^*). **(Q, R)** Expressing **(Q)** *Pdk^RNAi^* (*domeMeso-Gal4,UAS-GFP;UAS-Pdk^RNAi^*) or **(R)** *Pdha^RNAi^* (*domeMeso-Gal4,UAS-GFP;UAS-Pdha^RNAi^*), specifically in the progenitor cells, does not reveal any dramatic change in GSH levels. Compare to control **(G)**. **(S, S’)** Expressing **(S, S’)** *Pdha^RNAi^* (*domeMeso-Gal4,UAS-GFP;UAS-Pdha^RNAi^*) specifically in the progenitor cells, does not show any change in their cysteine levels. Compare to control **(D)**. **(T)** Relative steady state levels (fold change, f.c.) of glutamate in *domeMeso>GFP/+* (control, N=7, n=23)*, domeMeso>GFP/Gat^RNAi^* (N=4, n=14, p=0.8275) and *domeMeso>GFP/Ssadh^RNAi^* (N=5, n=17, p=0.7971). **(U)** Relative steady state levels (fold change, f.c.) of glycine in *domeMeso>GFP/+* (control, N=7, n=23)*, domeMeso>GFP/Gat^RNAi^* (N=4, n=14, p=0.5428) and *domeMeso>GFP/Ssadh^RNAi^* (N=5, n=17, p=0.9311). **(V)** Quantification of blood-progenitor cysteine levels (fold change, f.c.) in *domeMeso>GFP/+* (control, n=50)*, domeMeso>GFP/Gat^RNAi^* (n=26, p=0.0079), *domeMeso>GFP/Pdha^RNAi^;Gat^RNAi^* (n=30, p<0.0001 in comparison to *Gat^RNAi^*) and *domeMeso>GFP/Pdha^RNAi^* (n=26, p=0.5386). **(W)** Quantification of blood-progenitor GSH levels (fold change, f.c.) in *domeMeso>GFP/+* (control, n=59)*, domeMeso>GFP/Gat^RNAi^* (n=28, p<0.0001), *domeMeso>GFP/Pdha^RNAi^;Gat^RNAi^* (n=41, p=0.0001 in comparison to *Gat^RNAi^*), *domeMeso>GFP/Pdk^RNAi^* (n=33, p=0.5488) and *domeMeso>GFP/Pdha^RNAi^* (n=51, p=0.0301).

**Figure S3.**
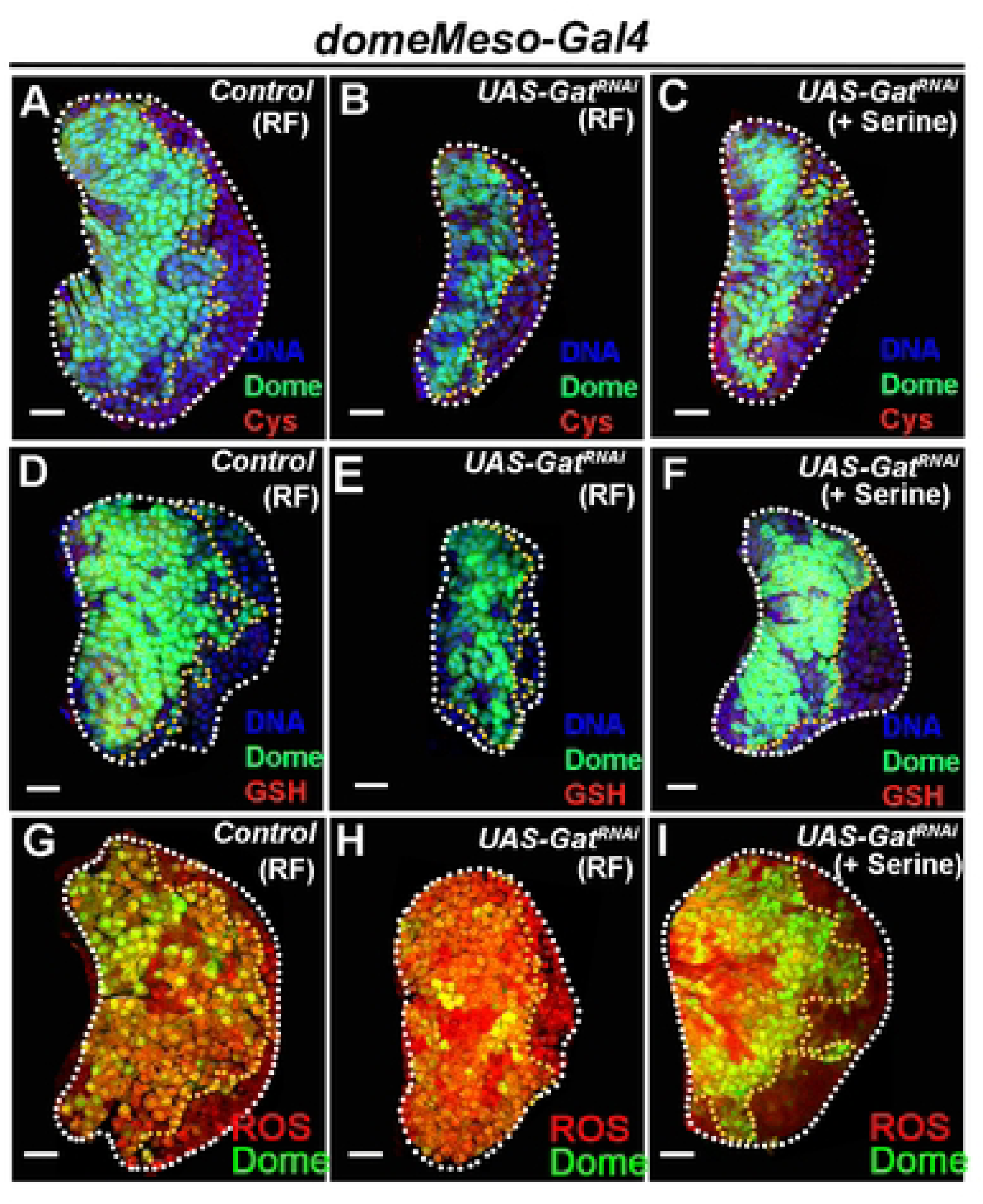
GABA catabolism controls progenitor cysteine synthesis via regulating de novo serine synthesis and maintain ROS homeostasis in the progenitor cells. RF is regular food, and serine is serine supplemented food. Scale bar: 20µm. DNA is stained with DAPI in blue, dome marks the progenitor cells in green. White border demarcates the lymph gland lobe and yellow border marks the dome positive area towards the left side. **(A-C)** Representative images showing cysteine (Cys, red) expression in lymph gland progenitor cells (area marked within the yellow dotted line) with merge of dome+ (green), DNA (blue) and cysteine (red) from different genetic backgrounds. **(A)** Control (RF, *domeMeso-Gal4,UAS-GFP/+*) lymph gland showing relatively uniform cysteine levels in all cells of the lymph gland including progenitor-cells (area demarcated within the yellow border). While, expressing **(B)** *Gat^RNAi^* (RF, *domeMeso-Gal4,UAS-GFP;UAS-Gat^RNAi^*) in the progenitor cells leads to reduction in cysteine levels as compared to control **(A)**, supplementing this genetic condition with **(C)** serine (Serine, *domeMeso-Gal4,UAS-GFP;UAS-Gat^RNAi^*) recovers cysteine levels almost comparable to control **(A)**. For comparison, also refer to **(B)** *Gat^RNAi^* raised on regular food (RF). **(D-F)** Representative images showing GSH (red) expression in lymph gland progenitor cells (area marked within the yellow dotted line) with merge of dome+ (green), DNA (blue) and GSH (red) from different genetic backgrounds. **(D)** Control (RF, *domeMeso-Gal4,UAS-GFP/+*) lymph gland showing GSH levels in lymph gland progenitor-cells (area demarcated within the yellow border). While, expressing **(E)** *Gat^RNAi^* (RF, *domeMeso-Gal4,UAS-GFP;UAS-Gat^RNAi^*) in the progenitor cells leads to reduction in GSH levels as compared to control **(D)**, supplementing this genetic condition with **(F)** serine (Serine, *domeMeso-Gal4,UAS-GFP;UAS-Gat^RNAi^*) recovers GSH levels almost comparable to control **(D)**. For comparison, also refer to **(E)** *Gat^RNAi^* raised on regular food (RF). **(G-I)** Representative images showing ROS levels (area marked within the yellow dotted line) with merge of dome+ (green) and ROS (red) from different genetic backgrounds. **(G)** control (RF, *domeMeso-Gal4,UAS-GFP/+*) lymph gland showing ROS levels in lymph gland progenitor-cells (area demarcated within the yellow border). While, expressing **(H)** *Gat^RNAi^* (RF, *domeMeso-Gal4,UAS-GFP;UAS-Gat^RNAi^*) leads to increase in progenitor ROS as compared to control **(G)**, supplementing this genetic condition with **(I)** serine (Serine, *domeMeso-Gal4,UAS-GFP;UAS-Gat^RNAi^*) recovers the increased ROS levels. For comparison, also refer to **(H)** *Gat^RNAi^* raised on regular food (RF).

